# Hepatitis B mother-to-child transmission in the Eastern Region of Ghana: A cross-sectional pilot study

**DOI:** 10.1101/402479

**Authors:** Thomas Hambridge, Yvonne Nartey, Amoako Duah, Amelie Plymoth

**Affiliations:** Department of Medical Epidemiology and Biostatistics (MEB), Karolinska Institutet, Stockholm, Stockholm County, Sweden; Department of Medicine and Therapeutics, Cape Coast Teaching Hospital, Cape Coast, Central Region, Ghana; Department of Medicine, St. Dominic’s Hospital, Akwatia, Eastern Region, Ghana

## Abstract

**Background:** Hepatitis B is a major health concern in Ghana, where prevalence of the virus remains high and most chronic patients are infected during childhood or at birth. There are several factors which can influence transmission risk from an infected mother to her infant, such as the presence of viral markers, the viral load and the use of prophylactic interventions. It is therefore important to determine the prevalence and main factors associated with mother-to-child transmission of hepatitis B in the context of Ghana.

**Materials and methods:** In this cross-sectional pilot study, hepatitis B testing was performed on infants born to infected mothers at a single site in the Eastern Region of Ghana. Test results were matched to a questionnaire which consisted of variables related to pregnancy and birth conditions. This was primarily a descriptive study to determine the prevalence of hepatitis B mother-to-child transmission as well as the preventive interventions and diagnostic methods used. The study variables were also analysed independently using Fisher’s exact test, while mother’s age at the time of delivery was assessed using univariate logistic regression.

**Results:** A total of 51 cases were included in the study and three (5.9%) of the infants tested positive. No significant association was observed between mother’s age and mother-to-child transmission (OR: 1.077, 95% CI: 0.828 – 1.403, p=0.58). A majority of infants received the standard hepatitis B vaccination schedule (96.1%) while two-thirds received the birth dose. There was no significant association observed between the clinical interventions reported in the study and mother-to-child transmission. Testing for viral markers and the use of antiviral therapy during pregnancy were absent in the population and could not be reliably assessed.

**Conclusion:** There was a low prevalence of HBV mother-to-child transmission observed despite a clear absence of viral marker and viral load testing. It is recommended that viral profile analysis is performed for hepatitis B positive pregnancies to identify high risk cases.

## Introduction

Hepatitis B virus (HBV) remains a global health concern, with an estimated 2 billion people serologically positive worldwide [1]. According to a recent report from the World Health Organization (WHO), approximately 257 million individuals were chronically ill with the disease in 2015 [2]. Moreover, global mortality from hepatitis B is increasing, accounting for up to 1.2 million deaths per year [3]. In high prevalence areas, HBV infection is typically acquired during childhood or through perinatal transmission [4]. Mother-to-child transmission (MTCT) can refer to three separate modes of transmission: during pregnancy (intrauterine infection); during delivery; or postpartum horizontal transmission through breastfeeding or daily contact [5]. It is estimated that 90% of infected newborns will go on to develop chronic hepatitis B (CHB) and are at much higher risk of developing liver disease, including cirrhosis and hepatocellular carcinoma during their adulthood [6–9]. Thus, taking measures to stop MTCT is essential for reducing the burden of CHB.

In Ghana, chronic infection with HBV is considered highly endemic. In a recent systematic review of 30 studies across all 10 regions, Ofori-Asenso & Agyeman found that hepatitis B surface antigen (HBsAg) seropositivity was 12.3% [10]. These figures are more or less in line with a systematic review conducted by Schweitzer et al. which estimated that the prevalence of chronic HBV in Ghana was 12.9% [11]. Ghana introduced hepatitis B vaccination (DPT- Hib-HBV pentavalent vaccine) as part of the Expanded Programme on Immunization (EPI) in 2002, recommending administration of the vaccine at 6, 10 and 14 weeks. Additionally, WHO recommends administration of the HBV vaccine within 24 hours of birth. However, birth dose coverage remains low at 39% globally and Ghana is one of several countries yet to adopt this policy at a national level [12]. Moreover, The WHO Western Pacific Region highlighted in a recent report that post-vaccination serologic testing (PVST) for HBsAg and antibodies could be used to help evaluate infant seroprotection following immunization [13]. In addition to the birth dose vaccine, it is crucial to identify infants who are at higher risk of perinatal transmission to administer hepatitis B immune globulin (HBIG), which has previously been shown to be an effective form of post-exposure prophylaxis [14,15].

Hepatitis B e antigen (HBeAg) status and elevated viral load during pregnancy are established risk factors for perinatal transmission [16–18]. The rates of HBeAg seroconversion in chronically infected individuals, i.e. the clearance and development of antibodies against the antigen, increase with age [9]. This means that mother’s age during gestation could also be an important risk factor for transmission. Although there has been extensive research into the factors influencing HBV mother-to-child transmission, there has been limited focus on the context of West Africa and specifically Ghana. This is particularly relevant given the high prevalence of HBV genotype E in the region, a genetic variant of the virus which requires further investigation. Compared to other genetic variants of HBV, transmission of genotype E appears to be predominantly horizontal in endemic areas [19–21]. Moreover, HBeAg seroconversion likely occurs before child-bearing age in Ghana where genotype E is predominant and may result in lower rates of MTCT compared to other countries.

The main objective of this cross-sectional pilot study was to determine the prevalence of HBV mother-to-child transmission at a hospital in the Eastern Region of Ghana and to assess the clinical interventions employed. We also sought to identify the main factors associated with HBV mother-to-child transmission in this setting and to use these preliminary findings to develop a larger prospective cohort study to be conducted at multiple sites throughout Ghana.

## Materials and methods

### Study area

The study was conducted at St Dominic’s Hospital, Akwatia in the Eastern Region of Ghana. There are roughly 2200 babies born at St Dominic’s hospital each year and this site was selected based on the availability of accessible maternal health records and contact details for the mothers. The hospital is funded by donor and government contributions and does not have any specific protocols for handling cases of HBsAg positive pregnancies. They follow the national policy regarding birth dose vaccine and HBIG, both of which must be paid for out-of-pocket.

### Study population and subject selection

Convenience sampling of maternal health records was used to retrospectively identify mothers who were HBsAg positive during pregnancy within the specified study period, with their child aged between 6-24 months at the time of recruitment. At St. Dominic’s Hospital, HBsAg positive cases are recorded within the first days of visiting the antenatal care clinic and up until delivery. Using these records, a list was compiled which consisted of eligible mothers who gave birth within the specified time period. Cases were excluded where infants were outside of the study age range at the time of testing, as well as mothers who declined to provide consent.

### Sample analysis

Tests were performed using InTec HBsAg Rapid Test Strips of whole blood. According to the manufacturer, test strips have a sensitivity of 99.4% and specificity of 99.8%. Positive test results in infants were confirmed by laboratory analysis. Results from prospective HBsAg testing were matched to responses from a questionnaire completed by the mother, which included questions relating to the birth context and interventions.

The variables included in the questionnaire were selected based on evidence from previous studies in different settings. These included mother’s age at the time of delivery, the birth location, the birth type, the infant’s vaccination status, HBIG administration, mother’s HBeAg status, viral load measurements, PVST and whether antiviral treatment was administered during pregnancy. The questionnaire responses were a combination of recall by the mothers and information retrieved from government issued maternal and child health records.

### Data analyses

Questionnaire response and hepatitis B test data for all cases (n=51) was entered into Microsoft Excel 2007 and imported into Statistical Package for Social Sciences (SPSS) version 25 for analysis. Descriptive analysis was performed for all cases to assess the questionnaire response rates. This analysis was also intended to measure the scope of viral marker testing and interventions in HBsAg positive pregnancies and evaluate how cases are managed in the clinic. Categorical data was expressed here as a number and percentage.

Each of the responses collected from the questionnaire (aside from mother’s age and viral load) were categorical variables and analysed independently as unpaired data. The outcome used in the analysis of questionnaire responses was whether the child was HBsAg positive, i.e. whether mother-to-child transmission had occurred. The outcome was binary categorical. The data was cross-tabulated and analysed using Fisher’s exact test for independence. All p-values for this test were two-tailed with p<0.05 considered significant.

To investigate the effect of mother’s age at time of delivery on transmission risk, a univariate (unadjusted) logistic regression analysis was performed to determine whether there was an increased risk associated with incremental change in mother’s age. Odds ratio (OR) with 95% confidence intervals (CI) are presented. The p-values for the univariate logistic regression were two-tailed with p<0.05 considered significant. This pilot study was not sufficiently powered to perform multivariate logistic regression to adjust for covariates.

### Ethical considerations

The study was approved by the institutional ethical review board at St Dominic’s Hospital and local permission was obtained prior to recruitment. The details and reasons for the study were clearly explained in a language that was understood by participants. Information was provided including risks, benefits and any possible complications. Consenting mothers were asked to visit the clinic with their child for HBsAg testing and to complete the study questionnaire.

## Results

### General characteristics

A total of 51 cases were included in the study. Mother’s age in the sample population ranged from 23 to 40 years. The average age was 31.2 years and only two mothers recruited into the study were under the age of 25. All deliveries took place within a hospital setting with no home births observed. The percentage of caesarean sections was 37.3% and a majority of births were vaginal deliveries (62.7%) (Table 1)

**Table 1.** General characteristics of the mothers included in the study and the birth type, n=51

| Mother's age |  | Delivery |  |
| --- | --- | --- | --- |
| Parameter | Value (SD) | Birth type | Number of cases (%) |
| Mean age | 31.2 (4.6) | Home | 0 (0) |
| Median age | 31.5 | Hospital | 51 (100) |
| Minimum | 23 | Vaginal | 32 (62.7) |
| Maximum | 40 | Caesarean | 19 (37.3) |

### Clinical interventions

There was a high vaccination rate observed (96.1%), with only two of the infants receiving no HBV vaccine at all. Two-thirds of infants received the WHO recommended birth dose administered within the first 24 hours. Moreover, a majority of infants received the EPI three dose vaccine schedule at 6, 10 and 14 weeks of age in addition to the birth dose (59.2%). Despite the high vaccination rates, none of the infants underwent PVST to evaluate seroprotection status. More than half of the infants (60%) received single dose HBIG at birth to provide further protection. Only one mother received antiviral therapy during pregnancy and her HBV DNA was measured, which was less than 20 IU/ml. There were no HBeAg tests performed during pregnancy. However, HBeAg was tested on three occasions post-delivery, with all three tests returning positive results (Tables 2 and 3).

**Table 2.** Proportion of study participants receiving vaccination and prophylactic interventions

| Clinical interventions |  |
| --- | --- |
| Intervention | Number of cases (%) |
| **Vaccinated** |  |
| Yes | 49 (96.1) |
| No | 2 (3.9) |
| **Vaccine birth dose\*** |  |
| Yes | 32 (66.7) |
| No | 16 (33.3) |
| **Vaccine type** |  |
| Monovalent | 3 (6.1) |
| Pentavalent | 17 (34.7) |
| Both | 29 (59.2) |
| **PVST** |  |
| Yes | 0 (0) |
| No | 49 (100) |
| **HBIG administered<sup>†</sup>** |  |
| Yes | 30 (60.0) |
| No | 20 (40.0) |
| **Antiviral therapy** |  |
| Yes | 1 (2.0) |
| No | 50 (98.0) |

**Table 3.** Hepatitis B viral marker testing rates and results

| Viral marker testing |  |
| --- | --- |
| Testing | Number of Cases (%) |
| **HBeAg tested<sup>\*†</sup>** |  |
| Yes | 3 (6.0) |
| No | 47 (94.0) |
| **HBeAg result\*** |  |
| Positive | 3 (100) |
| Negative | 0 (0) |
| **HBV DNA tested** |  |
| Yes | 1 (2.0) |
| No | 50 (98.0) |
| **HBV DNA result** |  |
| <20 IU/ml | 1 (100) |

\*HBeAg testing performed after delivery
<sup>†</sup> one case uncertain whether HBeAg testing was performed

**Table 4.** Mother’s age at time of delivery in relation to HBV mother-to-child transmission

| Mother's age |  |  |
| --- | --- | --- |
| Odds Ratio | 95% CI | p-value |
| 1.077 | (0.828 – 1.403) | 0.58 |

**Table 5.**
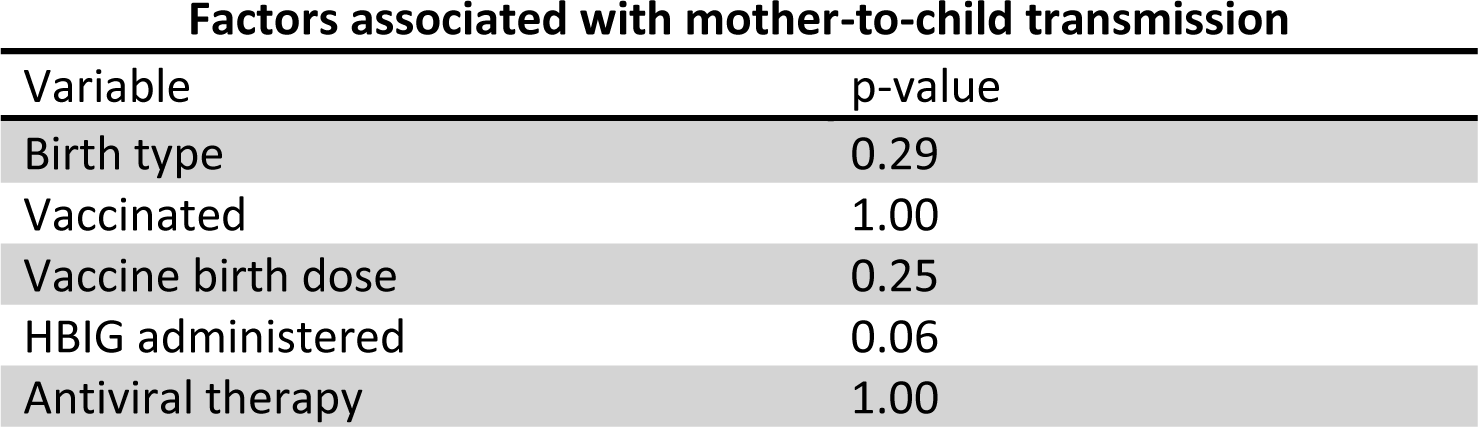
Factors related to HBV mother-to-child transmission

### Factors associated with HBV mother-to-child transmission

To investigate mother’s age at the time of delivery as a potential factor for transmission, a univariate logistic regression was performed with no covariates included in the model. This showed no significant association (OR: 1.077, 95% CI: 0.828 – 1.403, p=0.58). The remaining binary study variables were analysed using Fisher’s exact test of independence. Prevention of MTCT through administration of HBIG at birth was not significant (p=0.06), despite no transmission observed in any of the 30 cases that received the prophylaxis. None of the other study variables, including birth type, any vaccination, birth dose vaccine or antiviral therapy during pregnancy demonstrated significance using this statistical method. Data collected from the study questionnaire was insufficient to test for an association between HBeAg status or DNA viral load and risk of transmission.

## Discussion

This formative study aimed to determine the prevalence of HBV mother-to-child transmission at a single site in the Eastern Region of Ghana as a pre-trial for a larger study. The main factors associated with MTCT were investigated quantitatively, while the responses from the questionnaires were used to evaluate how HBsAg positive pregnancies were managed in the clinic. Given that the estimated national prevalence of individuals with CHB is approximately 12.3% in Ghana [10], the finding that only three of the 51 infants were HBsAg positive in this study was somewhat lower than anticipated. This is an important observation and appears to align more closely with the 8.3% HBV maternofetal transmission rate in chronic carriers observed by Candotti et al. in Kumasi [22] and the 8.7% seroprevalence reported by Amidu et al. in the nearby Ashanti region [23]. In most cases, when an adult is exposed to HBV, the virus will be cleared completely with no long-term manifestations. Conversely, a majority of those exposed to the virus during childhood when the immune system is immature will go on to develop lifelong infection [17]. If the prevalence of HBV mother-to-child transmission is in fact lower than expected in Ghana, this may support the argument that early horizontal transmission through direct contact is a major source of newly acquired chronic HBV infections in West Africa [22,24,25]. It should be highlighted here that we used a maximum inclusion age limit of 24 months to minimise false negatives while maintaining the assumption that any positive tests observed were the result of MTCT rather than early horizontal transmission. This assumption may therefore be a limitation of the study.

Although this pilot study was unable to demonstrate significant protection against HBV mother-to-child transmission with HBIG (p=0.06), this intervention has been previously shown to be an effective form of prevention [14,15]. The issue with the single-dose HBIG is that it is an expensive form of post-exposure prophylaxis and is not covered by the National Health Insurance Scheme (NHIS) in Ghana. There was no also no significant association observed using univariate logistic regression of mother’s age. There is rationale to explore this further with a larger sample size, as younger women are more likely to be in the HBeAg positive and high viral load phase known as ’immune tolerance’ [1]. It has been proposed that HBeAg seroconversion of HBV genotype E occurs at a much younger age than other genotypes, although this may still be an important factor to consider in younger women of childbearing age who have not yet seroconverted [26].

The vaccination rates observed at St. Dominic’s Hospital were encouraging, at 96.1% in total. A further 66.7% of all infants received the WHO recommended birth dose and only two received no HBV vaccine at all. The study was unable to demonstrate a significant association between vaccination and MTCT. Although speculative given the limited sample size, there is reason to hypothesise that sustained seroprotection may play a role in transmission risk, particularly with a potential lack of cross protection towards the endemic HBV genotype E [17,27]. This is currently an area of research interest in Ghana as vaccination uptake continues to increase and should be further investigated in the follow-up phase. It could also put a greater importance on PVST to ensure non-responders receive full vaccine protection. None of the infants in this study underwent PVST to evaluate seroprotection status.

One of the most important observations in this study was the absence of HBeAg and viral load testing. HBV viral profile testing can be performed locally to determine HBeAg status of infected mothers and may be used as a surrogate marker for high viral load. It has been consistently demonstrated in previous studies conducted in different settings that a positive HBeAg status and high viral load during pregnancy are strongly associated with elevated risk of MTCT [16–18]. In fact, of the three cases that performed HBeAg testing after delivery, each was positive for this marker and MTCT was observed in two of the three cases. Although these factors could not be reliably assessed here, the fact that no HBeAg test was performed and only one viral load was measured prior to delivery is a finding in itself. Moreover, they could represent key stages in the pregnancy and perinatal period which require further intervention to reduce the occurrence of MTCT in Ghana.

## Conclusion

There was a low prevalence of HBV mother-to-child transmission observed despite a clear absence of viral marker testing and subsequent intervention. Nevertheless, it is strongly recommended that positive HBsAg tests in Ghana are followed up with a viral profile analysis as an effective method of identifying high risk cases.

## Acknowledgements

We wish to express a special thanks to the staff at St. Dominic’s Hospital for their warm hospitality while conducting our research there and to all of the families who participated. We are also grateful to Professor Lewis Roberts at the Mayo Clinic and to the wider research team for all of their support and contributions.

## Author’s contributions

Study design: TH YN AD AP. Enrolled/recruited participants: AD. Performed the study: TH AD. Analysed the data: TH YN. Contributed materials and analysis support: AD AP. Wrote the paper: TH YN AD AP.

